# Divergent density feedback control of migratory predator recovery following sex-biased perturbations

**DOI:** 10.1101/828244

**Authors:** Daisuke Goto, Martin J. Hamel, Mark A. Pegg, Jeremy J. Hammen, Matthew L. Rugg, Valery E. Forbes

## Abstract

Uncertainty in risks posed by emerging stressors such as synthetic hormones impedes conservation efforts for threatened vertebrate populations. Synthetic hormones often induce sex-biased perturbations in exposed animals by disrupting gonad development and early life-history stage transitions, potentially diminishing per capita reproductive output of depleted populations and, in turn, being manifest as Allee effects. We use a spatially-explicit biophysical model to evaluate how sex-biased perturbation in life history traits of individuals (maternal investment in egg production and male-skewed sex allocation in offspring) modulates density feedback control of year class strength and recovery trajectories of a long-lived, migratory fish–shovelnose sturgeon (*Scaphirhynchus platorynchus*)–under spatially and temporally dynamic synthetic androgen exposure and habitat conditions. Simulations show that reduced efficiency of maternal investment in gonad development prolonged maturation time, increased the probability of skipped spawning, and, in turn, gradually shrunk spawner abundance, weakening year class strength. However, positive density feedback quickly disappeared (no Allee effect) once the exposure ceased. By contrast, responses to the demographic perturbation manifested as strong positive density feedback; an abrupt shift in year class strength and spawner abundance followed after more than two decades owing to persistent negative population growth (a strong Allee effect), reaching an alternate state without any sign of recovery. When combined with the energetic perturbation, positive density feedback of the demographic perturbation was dampened as extended maturation time reduced the frequency of producing male-biased offspring, allowing the population to maintain positive growth rate (a weak Allee effect) and gradually recover. The emergent patterns in long-term population projections illustrate that sex-biased perturbation in life history traits of individuals can interactively regulate the strength of density feedback in depleted populations such as *Scaphirhynchus* sturgeon to further diminish reproductive capacity and abundance, posing increasingly greater conservation challenges in chemically altered riverscapes.

## 1. INTRODUCTION

Human activities have uniquely shaped life history strategies of wild animal populations by imposing unnatural stress for centuries (Crain, Kroeker & Halpern 2008; Hendry, Farrugia & Kinnison 2008). The symptoms of human-induced stressors, in turn, manifest in life history traits varyingly among individuals (Marr *et al.* 2006) over time and space (Hope 2005; Maxwell *et al.* 2013). As stress becomes intensified, these sources of intraspecific variation in fitness may emerge as a directional shift in demographic rates (Wedekind 2017) as abundance declines. Positive density feedback following these demographic shifts may further diminish per capita reproductive capacity (a demographic Allee effect, Stephens, Sutherland & Freckleton 1999; Gregory *et al.* 2010), posing a threat to persistence of wild populations (Coltman *et al.* 2003).

One emerging stressor with such effects is synthetic chemicals, which have been increasingly detected in aquatic systems around the globe (Bernhardt, Rosi & Gessner 2017). A rising demand for human food production has increased agricultural use of synthetic chemicals (Tilman 1999; Tilman *et al.* 2001), posing a greater risk to freshwater vertebrates (Bernhardt, Rosi & Gessner 2017). Synthetic hormones–often used as growth-promoters for farmed animals–are especially potent and often induce sex-biased perturbation, impairing development and reproduction by disrupting energy allocation and maturation (del Carmen Alvarez & Fuiman 2005). Because maternal investment in egg production incurs greater energetic costs than somatic growth, females may more likely experience reproductive impairment than males (Arukwe & Goksøyr 1998). In pre-spawning females, even low-dose (often below detection) exposure to synthetic hormones can disrupt maturation processes such as vitellogenin production–a critical step in egg production (Arcand Hoy & Benson 1998; Nash *et al.* 2004), which can lead to skipped spawning in some fishes (Rideout, Rose & Burton 2005). Chemically induced inefficiency in egg production may, thus, act density-dependently especially in threatened and endangered populations as it reduces the number of females contributing to a reproductive event (known as a ‘mate-finding’ Allee effect, Gascoigne *et al.* 2009).

Synthetic hormones can also irreversibly perturb sex determination in exposed animals, skewing sex allocation in offspring (Mills & Chichester 2005). Resulting skewed adult sex ratio may manifest as a mate-finding Allee effect (Gascoigne *et al.* 2009) by impairing reproduction density-dependently (e.g., a decline in mating success), impeding recovery and posing extinction risk for depleted populations as abundance declines further. Such human-mediated modification in demographic traits has been reported in exploited populations with sex-biased mortality (Molnar *et al.* 2007; Grüebler *et al.* 2008). However, it remains unclear how positive density-dependent processes regulating fitness (also known as ‘component’ Allee effects, Stephens, Sutherland & Freckleton 1999) manifested from modified traits such as skewed sex ratio and delayed maturation act together to influence per capita reproductive output and population recovery in nature.

We here present a simulation study on how sex-biased, density-dependent perturbation in life history traits can emerge and modify recovery potential of a long-lived, migratory fish exposed to low-level synthetic androgens in a large river. Migratory animals are exposed to spatially and temporally dynamic flow-dependent exposure regimes as the chemicals move throughout networks of streams and rivers (O’Brien *et al.* 2016); variable exposure and subsequent symptoms among exposed individuals can obscure a causal link, if any, to population-level responses, delaying management actions (Peterson *et al.* 2003). We test varying exposure and effect scenarios on the Missouri River shovelnose sturgeon (*Scaphirhynchus platorynchus*, Fig. 1a) as an illustrative case study. Population sizes of *Scaphirhynchus* sturgeon have declined in many of their endemic habitats, and some are facing potential extirpation owing to extensive habitat modification (e.g., impoundments) (Bramblett & White 2001) that disrupt reproductive activities including seasonal spawning migration and aggregation. Some of the Missouri River populations have shown signs of unsuccessful spawning (DeLonay *et al.* 2009a) and year class failure owing to low larval fish survival (Hrabik *et al.* 2007)–a critical process for population persistence (DeLonay *et al.* 2009a). Because female *Scaphirhynchus* sturgeon mature later, spawn less frequently, and live longer than males (DeLonay *et al.* 2009a), female-biased disruption in reproductive traits likely regulate the strength of, if any, Allee effects. We use a spatially-explicit individual-based biophysical model that accounts for sex-specific life history traits to evaluate how sex-biased perturbation in energetics (reduced maternal investment in egg production) and demographics (male-skewed sex allocation in offspring)–two commonly reported symptoms of synthetic androgens–disrupts density-dependent reproductive processes to control year class success and population recovery under spatially and temporally dynamic exposure and habitat conditions.

**Figure 1.**
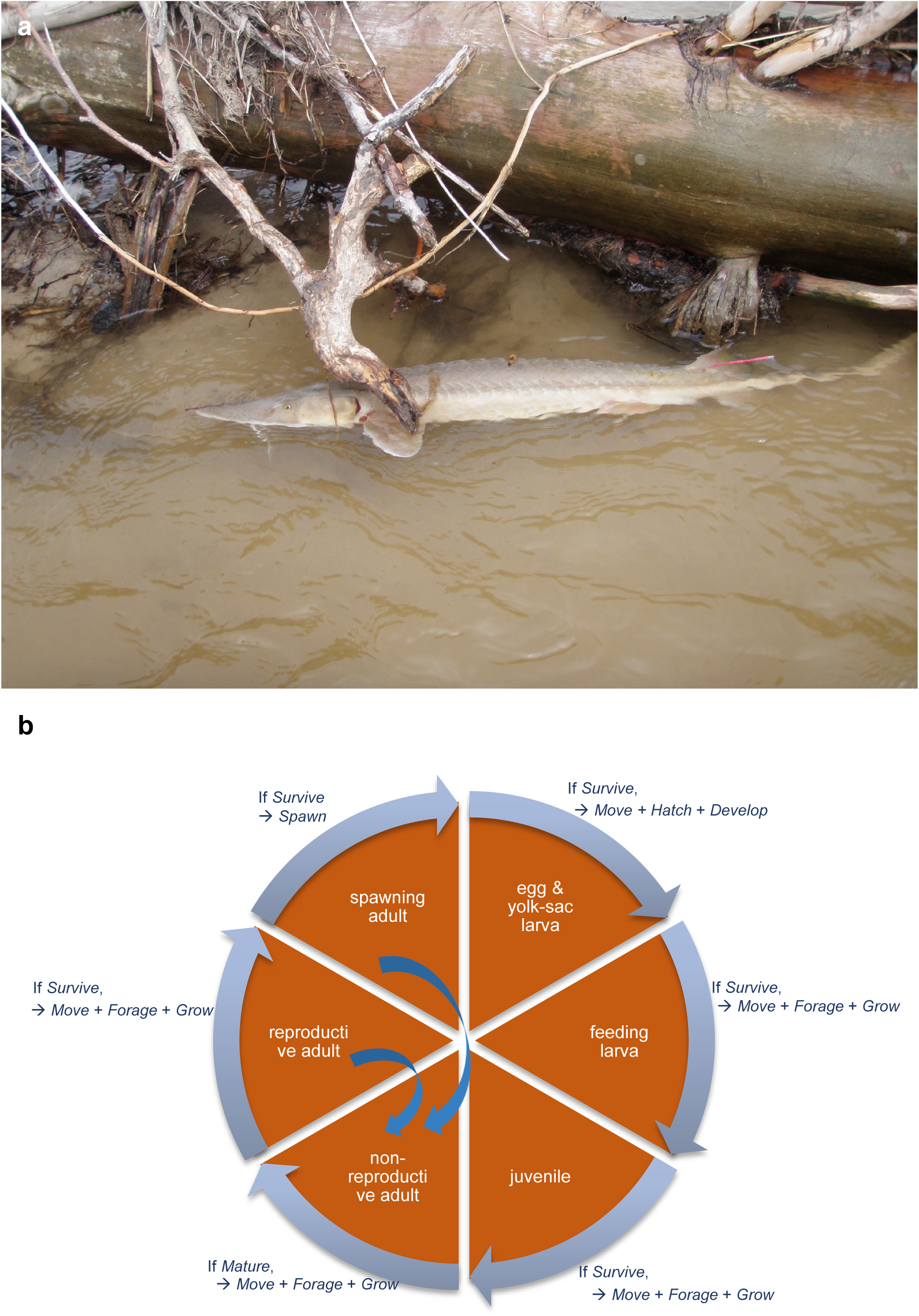
Shovelnose sturgeon *Scaphirhynchus platorynchus*. a) adult shovelnose sturgeon; b) a schematic of the spatially explicit individual-based biophysical model used in this study.

## 2. MATERIALS AND METHODS

### 2.1. Study system

The study system is the lower 162 km of the Platte River (Nebraska, USA), one of the largest tributaries of the Missouri River basin (Fig. 2 and Appendix 1: Fig. A1), providing spawning and nursing habitats essential for threatened populations such as *Scaphirhynchus* sturgeon (Peters & Parham 2008). Owing to its proximity to wastewater treatment plants and agricultural fields, however, the lower Platte River is also one of the impaired water bodies in the region (NDEQ 2004). Fish that inhabit this river and its tributaries, such as *Scaphirhynchus* sturgeon, are exposed to elevated concentrations of synthetic hormones widely used at concentrated cattle feedlots near the lower Platte River watershed (Huntzinger & Ellis 1993) and exhibit symptoms of endocrine disruption (e.g., intersexuality and atresia, DeLonay *et al.* 2009a).

**Figure 2.**
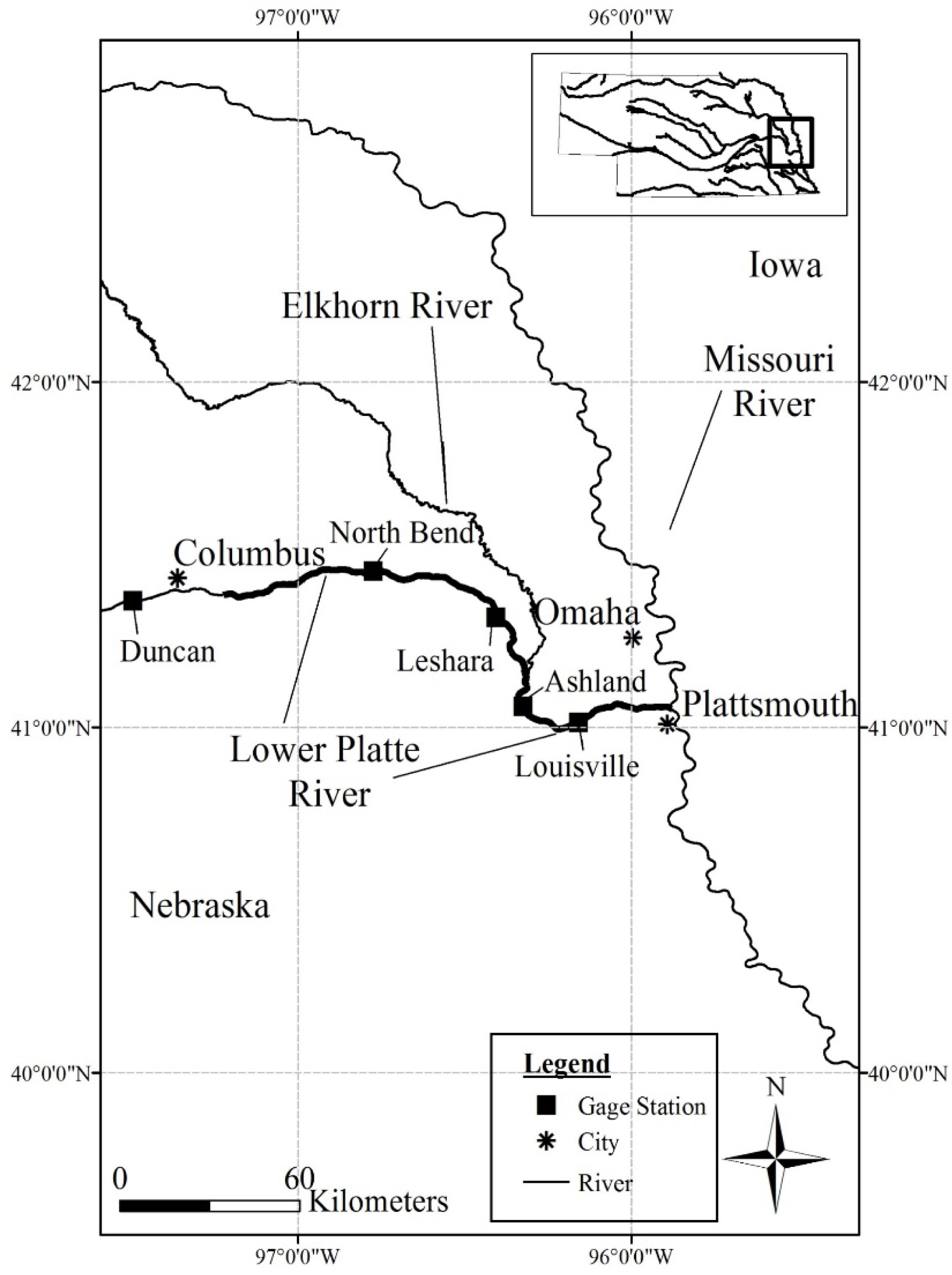
Map of the study system, lower Platte River, Nebraska, USA.

### 2.2. Sturgeon biophysical model

We use a spatially explicit individual-based biophysical model of shovelnose sturgoen previously developed for the lower Platte River popuatlion (Fig. 1b); the model is fully described in Goto *et al.* (2015) and Goto *et al.* (2018). The following briefly describes the model according to the Overview, Design concepts, and Details (ODD) protocol (Grimm *et al.* 2010).

#### Purpose

The model is designed to uncover mechanisms underlying early life-history processes modified by natural and human-induced stressors (e.g., synthetic chemicals), and their contributions to population persistence in a spatially and temporally dynamic environment.

#### Design concepts

1) *Basic principles*: The model primarily hinges on energetics principles to understand how maternal and environmental influences modulate year class variability through energy acquisition and allocation (Houde 1989); 2) *Emergence*: Demographic traits and spatial distribution of the population emerge from individual-level processes in response to habitat and indirect intraspecific competition. 3) *Sensing*: Sturgeon sense gradients of habitat conditions and prey density over neighboring cells; 4) *Interaction*: Sturgeon indirectly compete with each other for food in the same cell (density dependence is incorporated in directly *Foraging* and indirectly in *Growth*, Koch *et al.* 2009); 5) *Stochasticity*: Sturgeon’s abilities to sense habitat conditions, capture prey, decide when to spawn, and survive are stochastic because of inherent uncertainty and randomness; and 6) *Observation*: The model stores sturgeon state variables daily to compute demographic and life history trait values.

In this study, we evaluated sex-biased effects of synthetic androgen perturbation in maternal investment in gonad development and first-year sex allocation. The following briefly describes the essential features of the model relevant to this study.

#### Entities, state variables, and scales

The model simulates sturgeon population dynamics with two types of entities; 1) river and 2) sturgeon. The model river consists of 162 equally divided longitudinal rectangular segments (i.e., 1 river km each) with six state variables; daily, depth-averaged discharge (m^3^ · s^-1^), mean depth (gage height, m), channel width (m), benthic invertebrate prey density (g · m^-2^), water temperature (°C), and photoperiod (hrs). Discharge, depth, width, and prey density are spatially explicit, whereas temperature and photoperiod are spatially uniform. The model keeps track of 11 state variables, body length (mm), storage mass (g), structural mass (g), gonadal mass (g), the number of individuals in each super-individual, maturity status, reproductive status, physiological condition (see *Growth*), sex, age (years), and longitudinal location in the river (rkm), of the entire life cycle (Fig. 1b). Early life stages (fertilized eggs, yolk-sac larvae, and feeding larvae, and age-0 juveniles) are represented as super-individuals (Scheffer *et al.* 1995), whereas age 1+ juveniles and adults are represented as individuals. Each super-individual is a collection of individuals (“cohorts”) from the same mother.

#### Process overview and scheduling

The model simulates the population and updates state variables at discrete daily time steps over years and generations. The habitat conditions (including prey production) are first updated, and then the model evaluates actions by sturgeon in the following order; (i) *spawning*, (ii) *hatching and larval development*, (iii) *movement*, (iv) *foraging*, (v) *growth*, and (vi) *mortality* (Fig. 1b). The model updates state variables at the end of each action. At the beginning of each year, maturity is evaluated, and if mature sturgeon are healthy (see *Growth* below), they enter the reproductive cycle. Spawning normally occurs from late spring to early summer in the study system.

#### Submodels

The model consists of one benthic prey submodel and six sturgeon submodels. In this study, simulations were designed to evaluate the effects of synthetic androgen perturbations in vitellogenin production in adult females and sex allocation in offspring (see *Synthetic androgen effects* below for detailed description) through the following processes in the submodels:

a. *Benthic prey production* is computed using a mass-balance model of standing biomass, daily production, and net migration from neighboring cells as described by Rashleigh & Grossman (2005) with modifications described in Goto *et al.* (2018). Prey biomass density in each cell is initialized on January 1st of each simulation year with a mean of 18.8 (wet g · m^-2^) + 2.2RN, where RN is a random number drawn from the normal distribution (Whiles and Goldowitz, 2005).
b. *Embryonic and yolk-sac larval development* is computed using cumulative fractional temperature-dependent nonlinear functions (Goto *et al.* 2018) fitted to data reported by Wang, Binkowski & Doroshov (1985). Hatched larvae are assumed to use their yolk sac for ∼10 days (at 17–18 °C) before exogenous feeding.
c. *Foraging* is computed using a modified version of the Beddington–DeAngelis functional response model (Beddington 1975; DeAngelis, Goldstein & O’Neill 1975), a function of encounter rate, handling time, size-dependent prey selectivity, and prey and predator densities. The exact prey length is randomly assigned using a uniform distribution (between minimum and maximum of each size class) each day (Goto *et al.* 2018).
d. *Growth* is simulated by the bioenergetics model (Hanson *et al.* 1997) modified to represent the assimilated energy allocated to storage (reversible) and structure (irreversible) (Gurney *et al.* 2003). When structure is less than optimal storage, feeding larvae and juveniles allocate surplus net energy to storage first until optimal storage is reached (Goto *et al.* 2018). Growth in length is then calculated using modified length–mass functions, where length increases only when structural mass increases. When the assimilated energy is less than the energy expended through metabolism, mass is lost only from storage without any change in structure and length (Goto *et al.* 2018). Sturgeon mature based on sex-specific probability, which is length-dependent functions derived from field observations of the Platte River population (Goto *et al.* 2018). Reproductive adults further allocate energy to gonad incrementally between spawning events. Adults initiate the reproductive cycle only when their physiological condition is above a threshold based on relative storage mass, current storage/optimal storage (Goto *et al.* 2018). Non-reproductive adults allocate energy only to structure and storage. The amount of daily energy allocated to gonad (J · d^-1^) by healthy adults (based on their physiological condition) is computed by dividing expected gonad energy content on the spawning day by the minimal spawning interval (365 days for males and 1095 days for females) plus the earliest expected spawning day (day of the year). Expected gonad energy is computed by an allometric function derived by calibration for the Platte River population. The amount of daily energy allocated to gonad is further adjusted when the physiological condition is below the threshold (Goto *et al.* 2018). While the earliest expected spawning day occurs in late March (when daylight hours >12 hours), the realized spawning day is determined by discharge rate, temperature, and relative gonad mass.
e. *Spawning* is simulated as a stochastic event and occurs only when spawning cues–water temperature, river discharge rate, daylight hours, and gonad development (gonadosomatic index or GSI), concurrently meet the thresholds (Delonay *et al.* 2016). Female *Scaphirhynchus* sturgeon spawn at highly variable river discharge rates among Lower Missouri River tributaries (Delonay *et al.* 2016). Because our sturgeon model showed little sensitivity to this parameter (Goto *et al.* 2015), a minimum threshold is set at 141.6 m^3^ · s^-1^, which is expected to reach roughly 50% connectivity (migratory pathway connectedness) in the lower Platte River (Peters & Parham 2008) and directly influences the sturgeon movement and subsequently habitat conditions for other individual-level actions. A minimum GSI to initiate spawning is set at 0.18 for females and 0.03 for males based on field observations (Wildhaber *et al.* 2007a). Water temperature suitable for female *Scaphirhynchus* sturgeon spawning is set the range at 12.6–24.9°C based on field observations (DeLonay *et al.* 2009b). Fecundity is calculated by dividing gonad energy on the spawning day by single egg energy, assuming constant egg quality (45.72 J · egg^-1^, USDA 2001). After spawning, fish lose all gonad mass, and their physiological condition and total mass are updated; fish do not re-enter the reproductive cycle until the physiological condition meets the threshold. When either temperature or discharge rate does not meet the thresholds, fish skip spawning. We assume that when the physiological condition does not meet the threshold to remain in the reproductive cycle, gonadal energy is reabsorbed and reallocated to storage.
f. *Movement* is tracked in the longitudinal direction only. The planktonic life stage fish (eggs and yolk-sac larvae) move by passive drift with water currents (Braaten *et al.* 2008). Feeding larvae (post-settlement fish, Kynard, Henyey & Horgan 2002), juveniles, and adults move in the direction with probability of attaining higher habitat quality based on discharge rate (Goto *et al.* 2018) using the empirical models developed for the lower Platte River population (Peters & Parham 2008). Habitat suitability is based on relative areas of exposed sandbars, open water, and shallow sandbar complexes. The new realized location is then determined using fish swimming speed and water velocity (Goto *et al.* 2018); the maximum distance that fish can travel daily in this study is thus limited within three (above or below the current cell) neighboring cells (or 3.0 rkm).
g. *Mortality* results from thermal stress (eggs and yolk-sac larvae), predation (eggs, larvae, and juveniles), starvation, and recreational fishing (age 3+). We assume that a constant proportion of eggs and yolk-sac larvae die when water temperature rises above 24 °C (Kappenman, Webb & Greenwood 2013). Starvation mortality depends on relative storage mass; starvation mortality is evaluated only when a proportion of storage mass relative to total mass drops below the threshold (optimal storage, Goto *et al.* 2018). Fishing mortality was derived from annual creel surveys in the Platte River (Peters & Parham 2008). Realized predation, starvation, and fishing mortality are a stochastic event. Realized mortality of eggs and larvae is computed by drawing a random number from the binomial distribution with mortality rates as the probability and the number of individuals represented by each SI_yoy_ as the number of trials. For age 1+ fish, each fish dies when a random number drawn from a uniform distribution is below the probability set above.

#### Input data

Model inputs include water temperature (°C), mean gauge height (depth, m), channel width (m), discharge rate (m^3^ · s^-1^), and photoperiod (hrs). Water temperature, gauge height, channel width, and discharge rate (1995–2011, Appendix 1: Fig. A and Goto et al. 2018) are derived from the USGS National Water Information System (http://waterdata.usgs.gov/nwis). Further details on derivation of physical and hydrological input data are described in Goto *et al.*

(2018).

#### Initialization

We initialized simulations with 60 000 age 1+ individuals on January 1st. The initial location of each sturgeon was randomly assigned to a cell with non-extreme water velocity (≤0.6 m · s^-1^) and depth (≥0.5 m). The initial age structure follows a Poisson distribution with mean age of 7, which is truncated at a maximum age of 16 based on field observations (Hamel *et al.* 2015), and the initial sex ratio is 1:1. Sex is randomly assigned to each fish, assuming a binomial distribution. For age 1+ sturgeon, initial lengths-at-age were computed using the von Bertalanffy growth function (Ricker 1975) parameterized for the lower Platte River population (Goto *et al.* 2018). Once spawned, fertilized eggs were initialized as super-individuals; the number of super-individuals is the number of mothers, and the number of individuals in each super-individual is the total number of eggs spawned by each mother. After settlement, feeding larvae were initialized as an individual with an initial total length of 15.6 (mm) + 0.84RN (Braaten *et al.* 2008), where RN is a random number drawn from the normal distribution; these fish were subsequently simulated with individual-level processes (e.g., growth) and traits (e.g., body size).

### 2.3. Simulation scenarios for sex-biased purtubations by synthetic androgens

We ran 57-year (27-year exposure + 30-year recovery) simulations for the following four scenarios to assess population performance under two types of perturbation in life history traits of individuals induced by low-level synthetic androgen exposure, energetics and demorgraphics;

1. No sex-biased purtubation–baseline;
2. Synthetic androgens perturb maternal energetic investment in egg production by imposing reduction in vitellogenin production rate;
3. Synthetic androgens perturb sex ratio in offspring by imposing male-skewed sex allocation at recruitment; and
4. Synthetic androgens perturb both maternal investment and sex allocation. We assumed no direct mortality caused by synthetic androgen exposure. Further, we assumed that: a) synthetic androgens are released between May 1st and October 31st (a six-month growing season); b) androgens affect only fish located below the confluence with the Elkhorn River (∼51 rkm from Plattsmouth, NE, Fig 2), where a number of cattle feedlots are located; and c) because the androgen concentration varies with stream flow rate, the realized exposure level for individual fish is spatially and temporally dynamic, and is thus adjusted based on flow rate in a grid cell in which the given fish is located.

#### Energetic perturbation

We explicitly evaluated the mechanism underlying synthetic androgen-induced reduction in egg production by simulating the dose-dependent reduction of vitellogenin production in individual fish based on an exponential decay function (Ankley *et al.* 2003). Vitellogenin (VTG) concentration (mg · ml^-1^) is converted to relative gonad mass (GSI) using an empirically derived linear function based on field survey data on the Missouri River shovelnose sturgeon reported by Wildhaber *et al.* (2007b) (Appendix 1: Fig. A2) as GSI = 0.984VTG + 35.103 (*R*^*2*^ = 0.99). Reduction in daily energy allocation to gonad (g · d^-1^) in exposed females was then computed based on somatic mass on the given day.

Because we lack information on a quantitative relationship between synthetic androgen exposure and *Scaphirhynchus* sturgeon vitelogenin production, we tested the hypothetical scenarios of 15%, 25%, and 35% reduction in daily vitellogenin production (Appendix 1: Fig. A2) in fish that are located at the Elkhorn River–Platte River confluence; these reduction rates in vitelogenin production approximately corresspond to the responses in fish exposed to environmentally realistic, low level sysnthetic androgens (e.g., <∼15 ng · l^-1^ 17*β*-trenbolone, a synthetic androgen commonly used in cattle feedlots, Ankley *et al.* 2003). For fish that are located in cells below the confluence, the reduction rates were proportionally adjusted by the difference in flow rate (e.g., if flow rates at the confluence and in the cell where a fish is located are 100 and 120 m^3^ · s^-1^, respectively, the realized reduction rate for this fish is 12.5% under the 15% reduction scenario). Because the model sturgeon cannot detect low-level synthetic androgens in water, we assumed that their movement and energy acquisition (foraging) vary independently of synthetic androgens.

#### Demographic perturbation

We implemented synthetic androgen-induced skewness in sex allocation by proportionally reducing the probability of being female. We tested the hypothetical scenarios of 20%, 60%, and 80% reduction in probability of being females for post-hatch fish that remain in the cell at or below the Elkhorn River–Platte River confluence; these rates approximately corresspond to the responses in fish exposed to environmentally realistic concentrations (e.g., <∼15 ng · l^-1^ 17*β*-trenbolone, Morthorst, Holbech & Bjerregaard 2010). We assumed that the probability of being female incrementally declines by remaining in the exposure area each day during the annual six-month exposure period (i.e., 0.11%, 0.33%, and 0.44%, respectively). For fish in cells below the confluence, the probability was proportionally adjusted by the difference in flow rates (as described above).

For all the scenarios, we used hydrological and temperature inputs empirically derived from the lower Platte River field surveys during 1995–2011 (Goto *et al.* 2018), from which we randomly selected each simulation year; we assumed no systematic change or shift in hydrological and thermal conditions in this study. Further, we assumed no interannual change in turbidity, channel geomorphology, and photoperiod. We ran ten simulations for each exposure scenario and computed summary statistics and % deviations from the baseline simulations in demographic and life-history traits that contribute to per capita reproductive output; adult sex ratio, relative reproductive female number (the proportion of adult females in the reproductive cycle), reproductive female number, spawner biomass, and recruit number.

## 3. RESULTS

### 3.1. Baseline sturgeon population dynamics

Under the baseline conditions, the sturgeon population remained stable over 57 years (Fig. 3a) with positive density dependence in per capita population growth rate (Fig. 3b). The population maintained, on average, ∼32% of adults being female (Appendix 1: Fig A3a), adult females allocated, on average, ∼9.4 % of total mass to gonad while they were in the reproductive cycle (Appendix 1: Fig. A3b); ∼22% of reproductive females (Appendix 1: Fig. A3c) eventually spawned (= ∼15.8 metric ton spawner biomass, Appendix 1: Fig A3d), producing, on average, 23 million eggs (or ∼33000 eggs spawner^-1^) annually (Appendix 1: Fig. A3e, and Table S1 and S2). Most spawning and subsequent larval settlement occurred below the confluence of the Elkhorn River and the Platte River (112 rkm) in late April to early May (Table S1 and S2); on average, 0.11% of first-year fish survived (Table S1 and S2) with ∼7100 recruits per year (Appendix 1: Fig. A3f). Although the recruit numbers varied highly among years, a non-linear (hump-shaped) density-dependent relationship emerged between spawners and recruits (Appendix 1: Fig. A3g).

**Figure 3.**
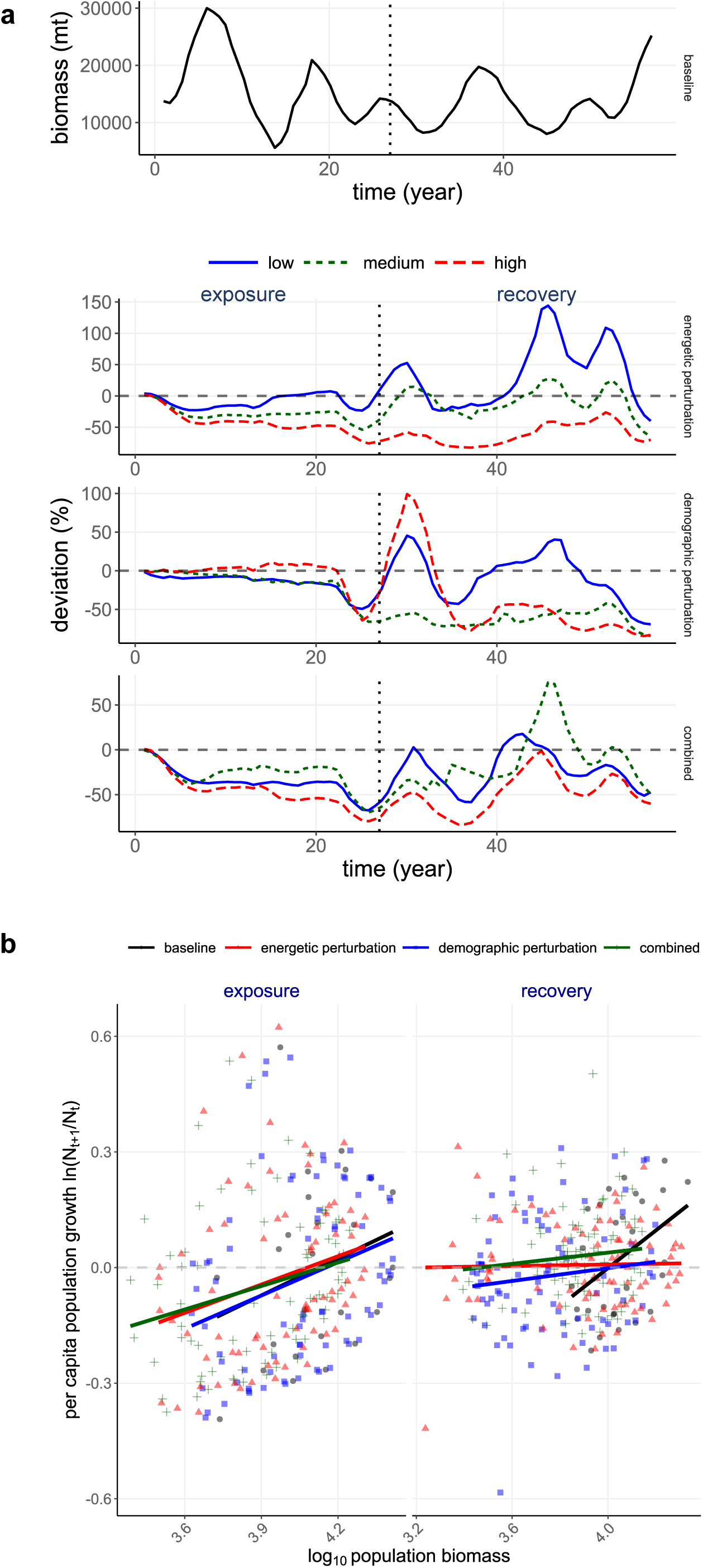
Population dynamics and density dependence in population growth of shovelnose sturgeon population. a) population projections during exposure and recovery under four synthetic androgen scenarios: a baseline scenario; an energetic perturbation (reduced maternal investment in gonad development) scenario with three exposure levels (shown as biomass relative to the baseline); a demographic perturbation (male-biased sex allocation in offspring) scenario; and a combined scenario of energetic and demographic perturbations. b) relationships between population biomass and per capita population growth rate, ln(*N*_*t*+*1*_/*N*_*t*_), during exposure (baseline: *y* = 0.32*x* – 1.3, *r* = 0.26; energetic perturbation: *y* = 0.24*x* – 1.0, *r* = 0.26; demographic perturbation: *y* = 0.29*x* – 1.2, *r* = 0.28; and combined: *y* = 0.20*x* – 0.84, *r* = 0.25) and recovery (baseline: *y* = 0.49*x* – 2.0, *r* = 0.43; energetic perturbation: *y* = 0.0098*x* – 0.031 *r* = 0.023; demographic perturbation: *y* = 0.083*x* – 0.33, *r* = 0.13; and combined: *y* = 0.072*x* – 0.25, *r* = 0.11).

### 3.2. Recovery from reduced vitellogenin production efficiency

Simulated reduction in vitellogenin production rate lowered reproductive capacities of the sturgeon population exposed to synthetic androgens (Fig. 4a). The proportion of energy allocated to gonad mass (10%) or storage mass (24%) in reproductive females under exposure did not differ from those under baseline (Table S1). However, the number of adult females in the reproductive cycle declined by 7.5–34% during exposure (Fig. 4a). Further, spawner biomass and recruit numbers declined, especially when exposed to moderate and high levels of the perturbation (6.8–25% and 23–38%, respectively, Fig. 4a), which were linearly correlated with relative changes in reproductive female abundance (Fig. 4b). Lowered reproductive capacities and year class resulted in an 8.6–43% reduction (relative to baseline) in population biomass (Fig. 3a) with positive density dependence in growth rate (Fig. 3b) during exposure.

**Figure 4.**
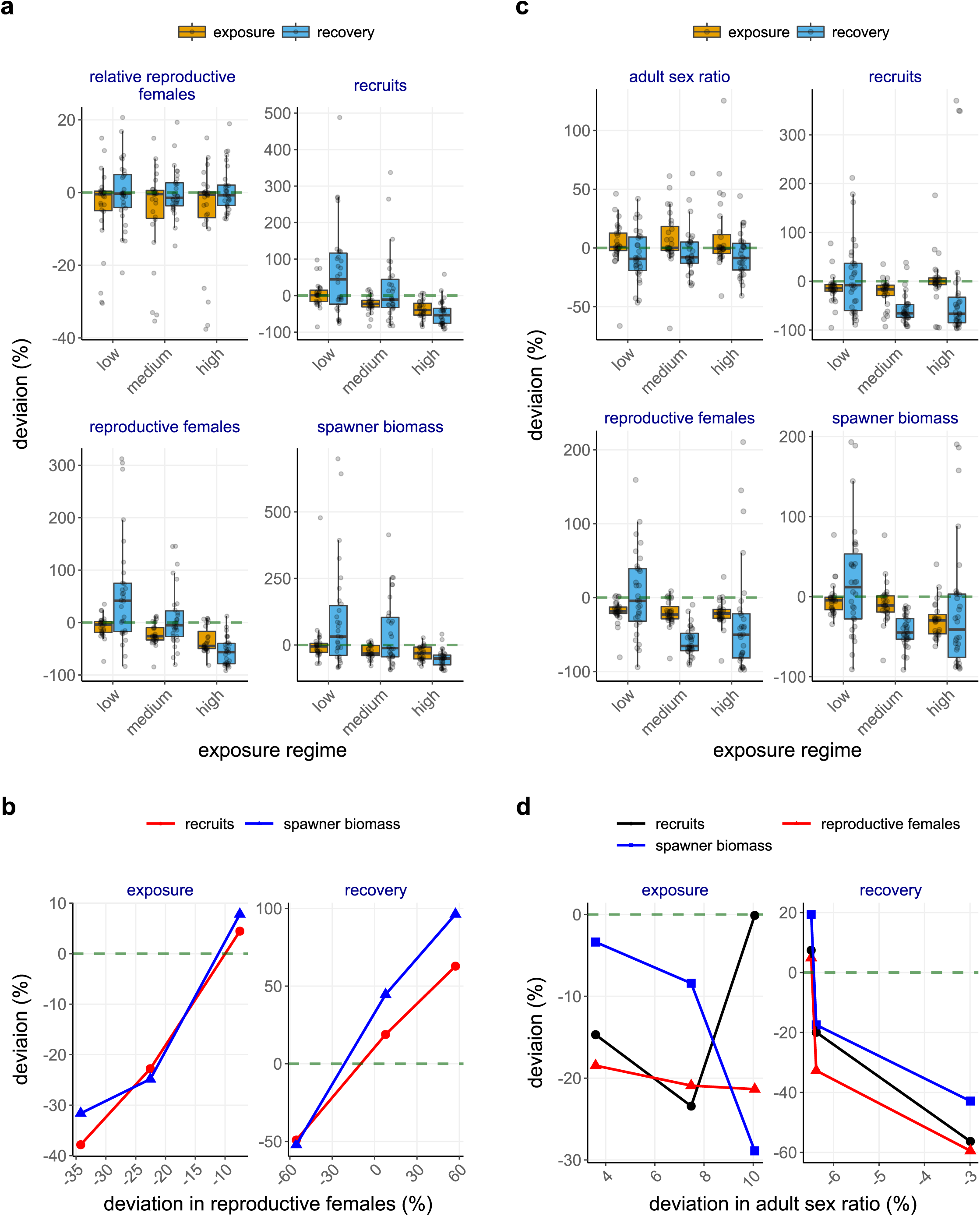
Deviation (%) compared to the baseline scenario in life history traits of shovelnose sturgeon populations. a) proportion of mature females in the reproductive cycle, recruit number, reproductive female number, and spawner biomass under energetic perturbation scenarios; b) relationship between relative reproductive female number and spawner biomass or recruit number during exposure and exposure under energetic perturbation scenarios; c) adult sex ratio, recruit number, reproductive female number, and spawner biomass under demographic perturbation scenarios, d) relationship between adult sex ratio and reproductive female number, spawner biomass, or recruit number during exposure and recovery under demographic perturbation scenarios. In a and c, black circles indicate annual mean values of life history traits. In b and d, data points show mean values from low, moderate, and high androgen exposure levels.

Reproductive females, spawners, and recruits of the population with moderate exposures returned to the baseline levels (Fig. 4a); similar linear relationships among the reproductive traits remained during recovery (Fig. 4b). Population size fully recovered (i.e., mean population size returns to the baseline level) within ∼five years (Fig. 3a) and density dependence in per capita growth rate virtually disappeared (Fig. 3b). However, the reproductive females, spawners, and recruits of the population with high exposure continued to decline (Fig. 4a), ultimately reducing mean population biomass by up to 83% before showing a sign of recovery (∼year 42, Fig. 3a).

### 3.3. Recovery from skewed recruit sex ratio

Sturgeon population biomass abruptly declined by up to 67% after ∼year 22 (Fig. 3a), and per capita growth rate continued to decline with biomass (Fig. 3b) under all scenarios of simulated male-skewed sex allocation during the first year of life. Skewed sex ratios in recruits were reflected in adults within ∼6 years with relative female numbers (the number of adult females entering the reproductive cycle) declining, on average, by 6.3–25% during exposure; mean reproductive female abundance declined by 19–21% (Fig. 4c). Subsequently, mean spawner biomass declined by up to 29% under high exposure (Fig. 4c). By contrast, mean recruit number declined under low and moderate exposure (by 15% and 23%, respectively), but differed little from the baseline population under high exposure (Fig. 4c). These changes in reproductive traits were, however, nonlinearly covaried with sex ratio skewness (Fig. 4d).

During recovery, the moderate and high exposure population biomass reached a stable state with less than 20% of the baseline population (Fig. 3a). Despite large year classes in early recovery years (year 31–33), up to more than 85% of adults were males under high exposure (Fig. 4c). On average, sex ratios under all the exposure scenarios remained skewed (by additional 9.3–16%) toward males with high interannual variation (Fig. 4c); sex ratio skewness did not show exposure-dependence and covary nonlinearly with reproductive traits (Fig. 4d). Although the low-exposure population fully recovered, the moderate and high exposure populations continued to have declining reproductive female abundance, spawner biomass, and recruit number (Fig. 4c), and, in turn, per capita growth rate continued to show positive density dependence and remained, on average, negative (Fig. 3b).

### 3.4. Recovery from combined effects of reduced vitellogenin production and skewed sex ratio

Concurrently imposing reduction in vitellogenin production and skewed sex ratios modulated the sturgeon population dynamics additively under low exposure, but antagonistically under moderate and high exposure. Population biomass declined within ∼5 years and remained stable and then declined further in the last five years (Fig. 3a). During exposure, adult sex ratio differed little under low exposure, but skewness toward males increased by 11% under medium and high exposure (Fig. 5a). Further, reproductive female abundance, spawner biomass, and recruit number declined by ∼31%, ∼23%, and ∼31% (respectively) under low and moderate exposures, and declined by 46%, 29%, and 47% (respectively) under high exposure (Fig. 5a), resulting in positive density dependence in per capita growth rate (Fig. 3b). Reproductive output (recruit number and spawner biomass) covaried nonlinearly with changes in relative reproductive number because of amplified responses under low exposure (Fig. 5b). By contrast, the relationships between reproductive traits (reproductive female number, spawner biomass, and recruit number) and adult sex ratio skewness abruptly shifted under high exposure (Fig. 5c).

**Figure 5.**
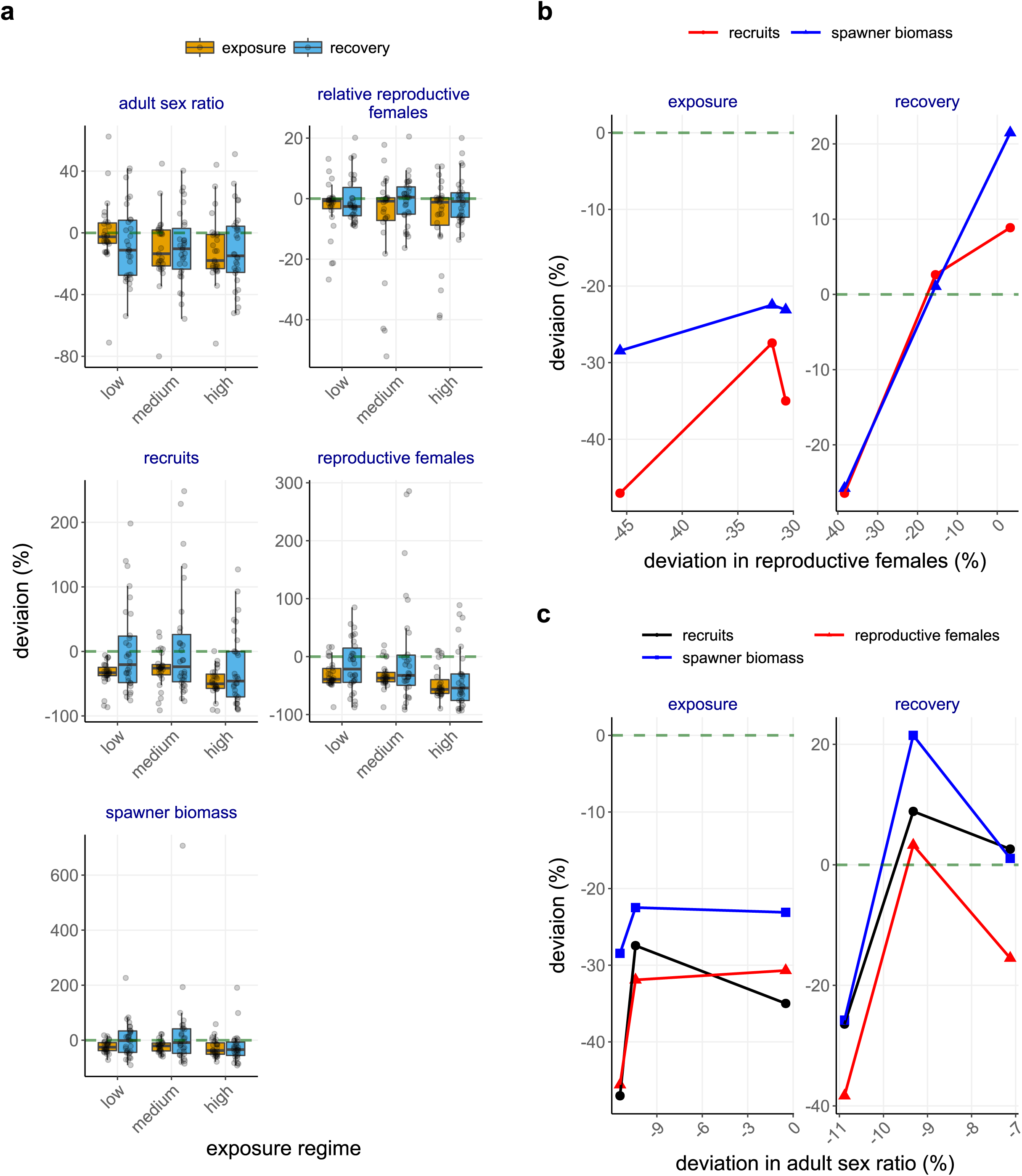
Deviation (%) compared to the baseline scenario in life history traits of shovelnose sturgeon populations under combined scenario of energetic and demographic perturbations. a) adult sex ratio, proportion of adult females in the reproductive cycle, recruit number, reproductive female number, and spawner biomass; b) relationship between relative reproductive female number and spawner biomass or recruit number during exposure and recovery, and c) relationship between adult sex ratio and reproductive female number, spawner biomass, or recruit number during exposure and recovery. In a, black circles indicate annual mean values of life history traits. In b and c, data points show mean values from low, moderate, and high androgen exposure levels.

Population biomass under low and moderate exposures recovered within ∼25 years, whereas population biomass under high exposure reached a stable state in ∼22 years at ∼44% lower than the baseline population (Fig. 3a). Although per capita growth rate continued to show positive density dependence, it remained, on average, positive during recovery (Fig. 3b). Adult sex ratios remained skewed toward males under all the exposure scenarios (Fig. 5a). Mean reproductive female number, spawner biomass, and recruit number (except for reproductive females under low exposure) recovered after low and moderate exposures but remained low under high exposure (Fig. 5a). The reproductive output covaried linearly with changes in relative adult female number (Fig. 5b); by contrast, the dampened responses in the reproductive traits during exposure also led to full recovery under medium as well as low (except for reproductive female number) exposures (Fig. 5b,c).

## 4. DISCUSSION

Simulations revealed plausible mechanisms involving chemically-induced, sex-biased perturbations in life history traits of long-lived, migratory vertebrates such as *Scaphirhynchus* sturgeon; these perturbations can emerge as vastly different response and recovery patterns– being manifest as differed strengths of demographic Allee effects (Hutchings 2015)–in a spatially and temporally dynamic environment. We find that lowered vitellogenin production rate prolongs maturation period and incurs skipped spawning (a relative decline in reproductive female abundance), and, in turn, reduces spawner abundance, ultimately weakening year class strength. Positive density feedback in per capita growth rate, however, disappeared (no Allee effect) during recovery. Further, the energetic perturbation operates as exposure-dependent processes–the higher the exposure level, the more severe the damage, thereby resulting in longer recovery time. By contrast, male-skewed sex allocation (an absolute decline in reproductive female abundance) emerges as delayed (by more than two decades) population-level responses, followed by an abrupt shift in year class strength and population trajectory under higher exposures, ultimately reaching an alternative state without any sign of recovery owing to persistent negative population growth (a strong Allee effect, Stephens, Sutherland & Freckleton 1999; Gascoigne *et al.* 2009). When combined with the energetic perturbation, however, impacts of the demographic perturbation are dampened as extended maturation time reduces the frequency of producing male-biased offspring, allowing the population to maintain positive growth rate (a weak Allee effect) and gradually recover. The emergent patterns in long-term population projections illustrate that sex-biased perturbation in life history traits of individuals can amplify or dampen Allee effects in depleted populations such as *Scaphirhynchus* sturgeon to further diminish reproductive capacity and abundance, posing conservation challenges in chemically altered riverscapes.

Sex-biased differences in energy allocation to gonadal maturation often determine life-history strategies such as foraging and mate selection, shaping sexual dimorphism in animal populations (Huse 1998; Rennie *et al.* 2008). Delayed gonad maturation in energetically perturbed females of our model sturgeon population prolongs the spawning cycle instead of reducing fecundity. Because sturgeon spend several years to build gonad energy prior to spawning (a capital spawner, McBride *et al.* 2015), slowed female gonad maturation and longer spawning cycle delay spawning timing. Although sturgeon are iteroparous, because of relatively infrequent spawning and long maturation time, delayed or skipped spawning events can reduce individual lifetime (in our simulations, some individuals spawned only once) and per capita annual reproductive output owing to fewer females contributing to spawning events (a reproductive Allee effect, Gascoigne *et al.* 2009). Inefficient energy use triggered by synthetic hormones in threatened animal populations with long maturation time can, thus, prolong population recovery time. Our simulations, however, suggest that the energetic perturbation also triggers plastic responses. Although reproductive and spawning females decline by up to 25%, the population eventually recovers partly because of limited positive density feedback in per capita growth rate following synthetic androgen exposure, indicating that reproductive capacity diminished during exposure can be restored if energy allocation to gonad resumes.

Delayed spawning events may, however, increase the probability of year class failures owing to environmental stochasticity. We find that exposed (energetically challenged) female sturgeon are more likely to abort the reproductive cycle, as they divert energy to meet metabolic needs when poor habitat conditions persist. Although evaluating energetic tradeoffs between migration and fecundity is beyond the scope of our study, animals with extensive spawning migration such as *Scaphirhynchus* sturgeon likely skip spawning or produce offspring with lower survival rate owing to reduced energy reserves (Glebe & Leggett 1981). Sufficient energy reserves in eggs are vital to larval development and survival; maternal investment in egg production determines the nutritional status of post-hatching larvae (Kjesbu *et al.* 1996). Our model assumes that individual eggs do not vary in energy content with synthetic androgen exposure; however, energetic tradeoffs in maternal investment in egg production and offspring survival can arise and vary greatly among taxa (Jönsson 1997; McBride *et al.* 2015) and thus need further investigation for *Scaphirhynchus* sturgeon.

Our study shows that recovery from demographic perturbation can pose greater conservation challenges than energetic perturbation. We find that chemically-induced, male-skewed adult sex ratio can incur a lasting mate-finding Allee effect (Gascoigne *et al.* 2009) in long-lived populations such as *Scaphirhynchus* sturgeon, severely diminishing their reproductive fitness and recovery potential. Although skewed sex ratio has been widely reported for a variety of vertebrate taxa (Donald 2007), causes of the skewness vary; it can result from sex-specific phenotypic differences in physiology, dispersal, and natural or human-induced mortality (Weimerskirch, Lallemand & Martin 2005; Goto 2009; Goto & Wallace 2010; Pen *et al.* 2010). In our baseline population, adult sex ratios are skewed toward males (∼1:2.2), primarily resulting from 1) a longer maturation time for females than males and 2) more energy investment to reproduction in females, resulting in higher probability of starvation mortality in poor environment years. Simulations show that further deviation (by 13–23%) in adult sex ratio skewness result in long-lasting threats to year class success and population viability. Consequences of skewed adult sex ratio may not immediately emerge for long-lived species (Holmes & York 2003; Lawson *et al.* 2010) such as *Scaphirhynchus* sturgeon. Once the skewness reaches a tipping point, however, it often drastically reduces recovery potential (Gascoigne *et al.* 2009) as it may stabilize in an alternative state (low abundance) facilitated by positive feedback (Van Nes & Scheffer 2007; Takimoto 2009; Dai *et al.* 2012). Although spawning aggregation formed by the Missouri River populations of *Scaphirhynchus* sturgeon may, to some extent, mitigate the component (mate-finding) Allee effect (Stephens, Sutherland & Freckleton 1999; Gascoigne *et al.* 2009), the threshold of this effect (female shortage) to avoid demographic shifts and facilitate recovery of these populations needs further investigation.

Finally, our study demonstrates that two or more component Allee effects can act interactively to dampen (or amplify) density feedback in per capita reproductive output and promote (or prevent) recovery in masculinized populations. Simulations show that by reallocating gonad energy to growth and postponing spawning events, surviving adult female sturgeon mitigate the strong Allee effect of skewed sex ratio by reducing the overall frequency of producing male-skewed offspring during exposure. In conservation planning for long-lived migratory species such as *Scaphirhynchus* sturgeon, thus, to prevent the shifts and minimize recovery cost, it is critical to identify early warnings that signal impending demographic shifts; the threshold of skewness in adult sex ratio that leads to demographic Allee effects can be a sensitive metric for assessing viability of threatened populations (Gascoigne *et al.* 2009; Ancona *et al.* 2017).

Long-term, low-level exposure to stressors such as synthetic hormones often obscures causative relationships (Windsor, Ormerod & Tyler 2018). Establishing ecologically relevant indicators such as the thresholds of Allee effects (where the shift occurs from positive to negative per capita growth) resulting from modified life history traits would guide the risk assessment of synthetic chemicals and habitat conservation planning for threatened populations. Further, given the time lags across biological hierarchy, continued biological monitoring of individual-level traits would help detect early signals that may indicate population-level changes such as the strength of Allee effects. Although our study was designed to evaluate consequences of low-level exposure, our findings also indicate larger implications of among-individual variation for population persistence and recovery of threatened and endangered species. Population models widely used in risk assessments often assume no among-individual variation in a population such as sex-, size-, or age- dependent life history traits (e.g., age at maturity). These models may not only gloss over individual variation in life history traits that may contribute to population viability, but also under- or over- estimate the strength of density feedback, and, in turn, the probability of extinction risk.

Large-river migratory populations such as *Scaphirhynchus* sturgeon experience multiple synthetic chemicals over agriculturally rich landscapes in regions such as the Midwestern United States (Welcomme 1995). Further confounding factors in conservation and management efforts can arise in hydrologically altered systems such as the Missouri River. Precipitation, for example, can regulate the amount of agrichemicals released into rivers. Assessing the likelihood of ecological risks of agrichemicals for long-lived species requires the development of trait-based predictive models using anticipated scenarios. With expected trends in rising climate variability, which can have considerable effects on river hydrology and water chemistry, these forecasting tools are critical for fisheries and water resource management in chemically altered ecosystems.

## AUTHORS’ CONTRIBUTIONS

DG, VEF, and MAP conceived the ideas; MJH, JJH and MLR collected and managed the data; DG led the writing of the manuscript. All authors contributed to revisions and gave final approval for publication.

## ACKNOWLEDGEMENTS

We greatly appreciate comments by Maxime Vaugeois (University of Minnesota) and anonymous reviewers on the earlier version of this manuscript. This project was partially funded by University of Nebraska–Lincoln and the Nebraska Game and Parks Commission (Project F-180-R).

## DATA AVAILABILITY STATEMENT

Model input data on water temperature, gauge height, channel width, and discharge rate used are publicly available at the United States Geological Survey (USGS) National Water Information System (http://waterdata.usgs.gov/nwis).

## FIGURE LEGENDS FOR APPENDIX 1

**Figure A1.**
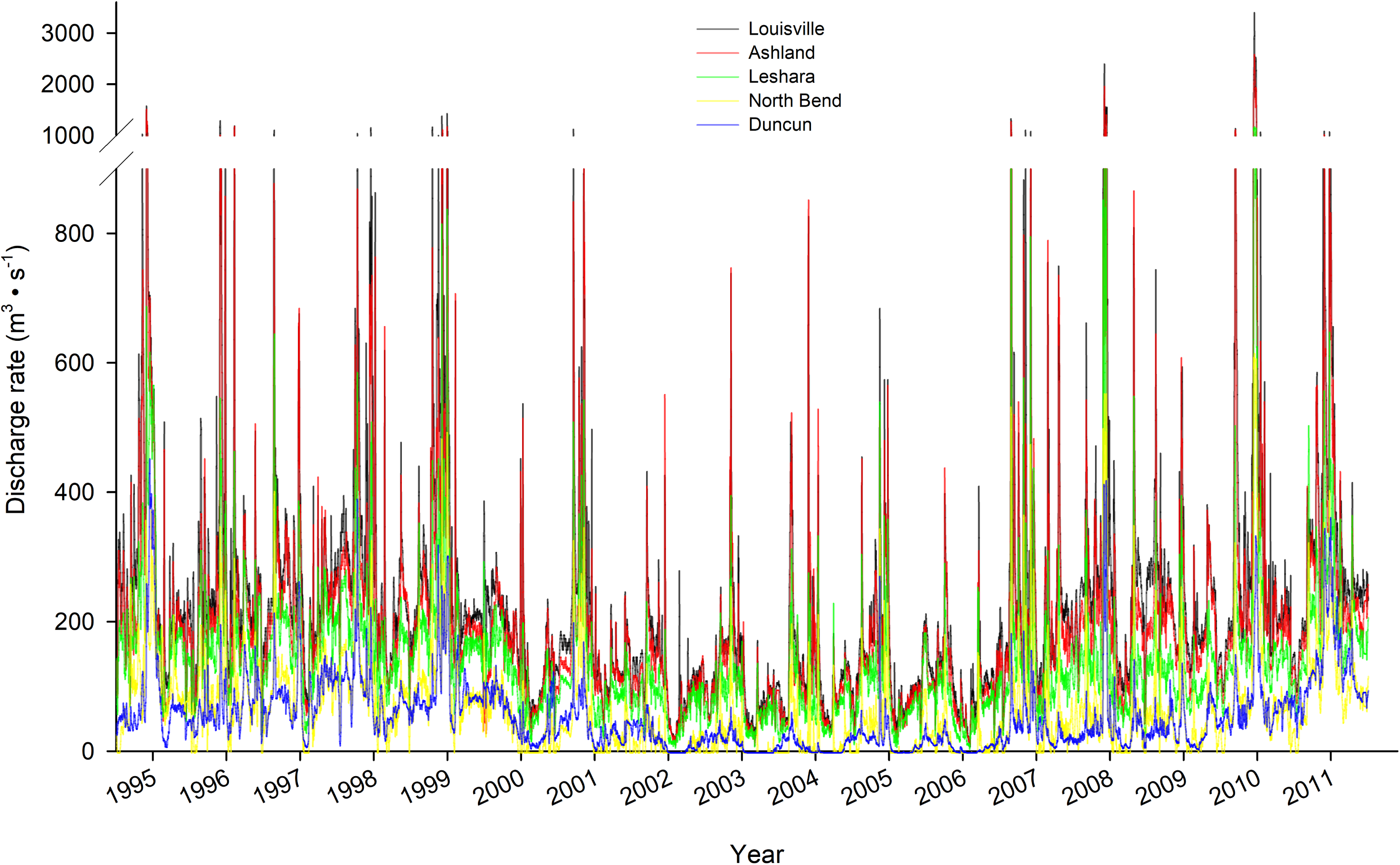
Daily mean river discharge rates measured at five USGS gaging stations (Duncan– station #06774000, North Bend–#06796000, Leshara–#06796500, Ashland–#06801000, and Louisville–#06805500) during 1995–2011.

**Figure A2.**
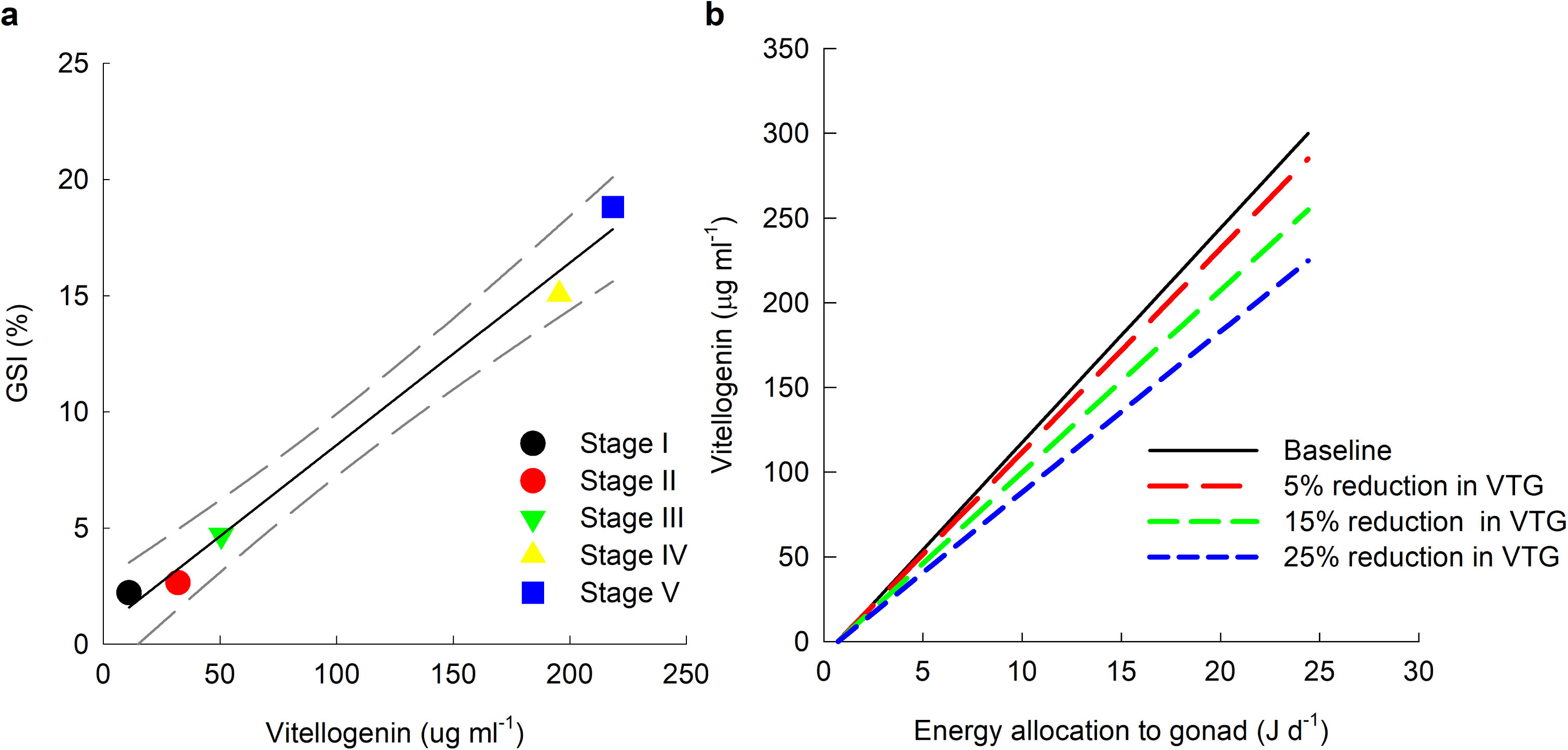
a) Relationship between energy allocation to female gonad development and vitellogenin computed from field estimates of vitellogenin concentrations and gonadosomatic index in pre-spawning female shovelnose sturgeon reported in Wildhaber et al. (2007); and b) simulation scenarios tested for the maternal investment in egg production (energetic perturbation). In a, Stage I, II, III, IV, and V represent immature, developing, vitellogenesis, pre-spawning, spawning, and spent (post-spawning) of the female gonad development (Wildhaber et al. 2007). In b, the baseline relationship is based on data reported in Wildhaber et al. (2007).

**Figure A3.**
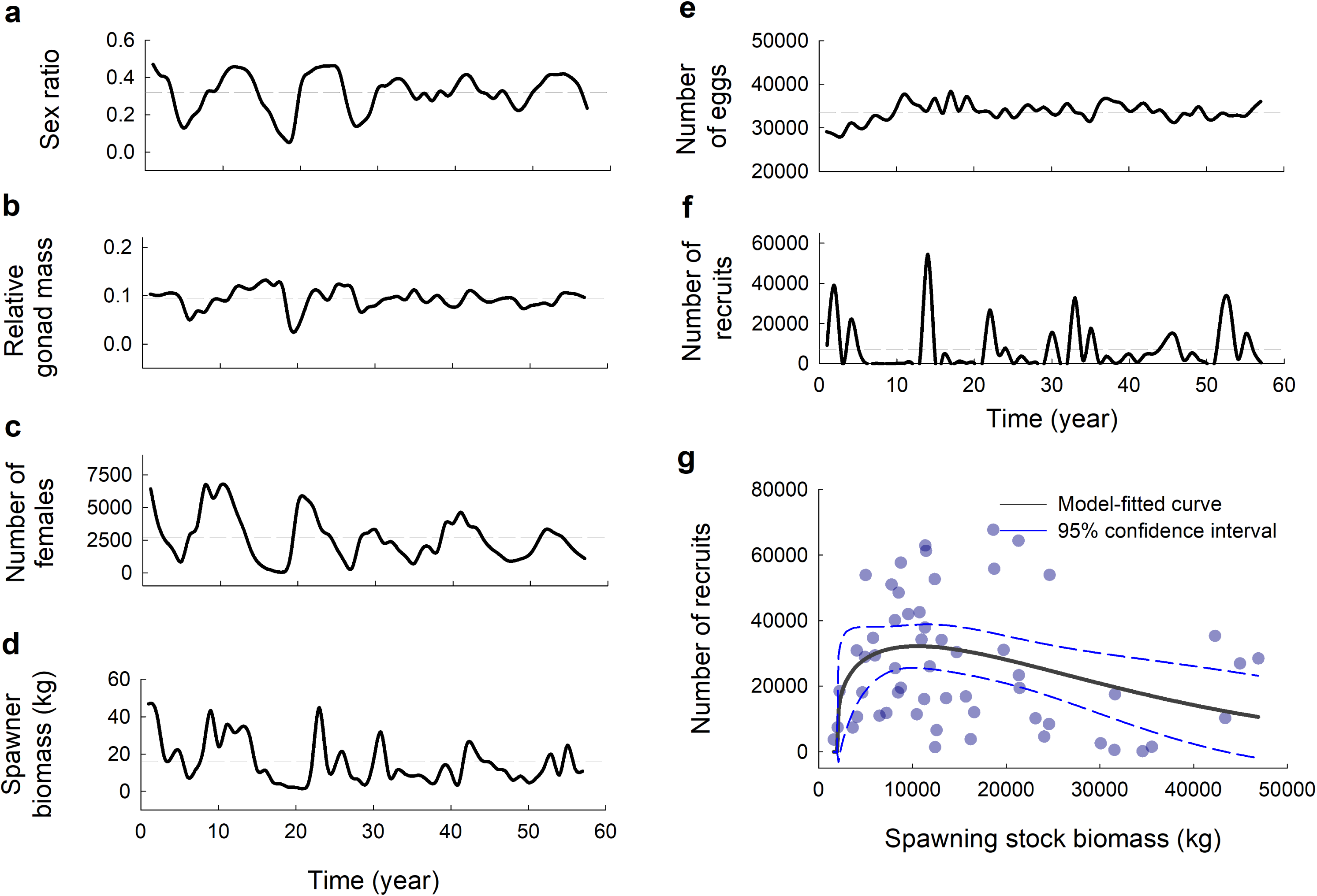
Temporal patterns of demographic and reproductive traits of simulated shovelnose sturgeon populations under the baseline scenario – a) adult sex ratio, b) female number, c) relative female gonad mass, d) fecundity, e) spawner biomass, f) recruit number, and g) a stock-recruit relation. The sturgeon model used for the baseline simulation has been calibrated and validated with field survey data on the lower Platte River population (Goto *et al.* 2015).

## Supporting Information

**Table S1.** Summary of exposure period (27 years) model outputs (mean ± sd) from the shovelnose sturgeon spatially explicit individual-based model simulations for energetic (reduced energy allocation to female gonad development) and demographic (pre-recruit sex ratio skewness) perturbation scenarios. Vt_low, Vt_mid, and Vt_high indicate vitellogenin reduction in adult females exposed to low-, mid-, and high- level synthetic androgen. SS_low, SS_mid, and SS_high indicate skewed sex ratio in larvae exposed to low-, mid-, and high- level synthetic androgen. Numbers indicates mean values computed across years.

| model output | baseline | Vt_low | Vt_mid | Vt_high | SS_low | SS_mid | SS_high | Vt + SS_low | Vt + SS_mid | Vt + SS_high |
| --- | --- | --- | --- | --- | --- | --- | --- | --- | --- | --- |
| **adult female** |  |  |  |  |  |  |  |  |  |  |
| sex ratio (proportion of female) | 0.31±0.15 | 0.32±0.13 | 0.32±0.13 | 0.32±0.13 | 0.29±0.14 | 0.27±0.13 | 0.24±0.11 | 0.31±0.12 | 0.27±0.11 | 0.27±0.12 |
| mean age of adult female | 6.88±0.56 | 6.86±0.52 | 6.85±0.52 | 6.86±0.52 | 6.90±0.57 | 6.92±0.56 | 6.89±0.55 | 6.88±0.53 | 6.87±0.54 | 6.86±0.52 |
| reproductive female abundance | 3119.0±2260.5 | 2819.4±1979.2 | 2505.4±1782.5 | 2002.4±1550.5 | 2643.6±2023.5 | 2532.6±1908.1 | 2475.2±1836.0 | 2138.0±1596.5 | 2092.3±1542.9 | 1648.4±1407.2 |
| mean relative storage mass of reproductive female | 0.24±0.04 | 0.24±0.04 | 0.24±0.04 | 0.24±0.04 | 0.24±0.04 | 0.24±0.04 | 0.24±0.04 | 0.24±0.04 | 0.24±0.04 | 0.24±0.03 |
| mean relative storage mass of non-reproductive | 0.25±0.02 | 0.24±0.05 | 0.24±0.02 | 0.25±0.02 | 0.24±0.04 | 0.25±0.03 | 0.24±0.04 | 0.24±0.03 | 0.24±0.02 | 0.24±0.03 |
| mean relative gonad mass of reproductive female | 0.10±0.03 | 0.10±0.03 | 0.10±0.03 | 0.10±0.03 | 0.10±0.03 | 0.10±0.03 | 0.10±0.03 | 0.10±0.03 | 0.10±0.03 | 0.10±0.03 |
| number of spawner | 701.6±708.3 | 714.3±670.4 | 680.1±670.7 | 467.1±572.5 | 604.4±646.7 | 600.0±653.5 | 577.6±645.5 | 500.8±593.8 | 480.0±577.6 | 406.2±558.2 |
| mean spawning location (rkm) | 103.9±12.9 | 103.7±13.2 | 100.1±13.9 | 99.0±13.9 | 103.4±11.3 | 103.9±15.7 | 104.0±15.2 | 102.2±17.1 | 100.4±15.5 | 124.8±14.0 |
| mean spawning time (day of year) | 117.6±12.1 | 116.0±10.6 | 115.3±10.6 | 113.7±10.6 | 117.3±14.3 | 118.5±12.6 | 118.5±11.6 | 115.6±11.0 | 114.1±12.7 | 113.5±11.5 |
| mean egg number per female | 33369.5±2697.1 | 33622.5±2570.8 | 33965.5±2571.0 | 34667.3±2882.0 | 32899.3±3189.0 | 33074.0±3232.3 | 33023.7±3173.8 | 33551.3±3217.0 | 33815.4±3145.3 | 34326.9±3423.3 |
| mean spawning interval (years) | 2.81±0.52 | 2.91±0.53 | 2.88±0.54 | 3.01±0.55 | 2.91±0.66 | 2.89±0.55 | 2.77±0.54 | 3.04±0.69 | 3.00±0.55 | 3.99±0.55 |
| **early life stages** |  |  |  |  |  |  |  |  |  |  |
| total egg number (millions) | 22.7±22.1 | 20.1±20.6 | 18.1±18.9 | 15.2±16.9 | 19.8±20.1 | 19.5±20.1 | 22.5±19.9 | 16.2±17.6 | 15.6±17.2 | 17.7±18.4 |
| mean larval settlement location (rkm) | 100.2±20.9 | 97.7±16.6 | 92.7±15.4 | 92.4±17.3 | 100.3±22.4 | 97.9±21.1 | 99.1±19.2 | 97.4±15.9 | 99.8±17.8 | 92.8±17.9 |
| mean first-year survival rate (%) | 0.11±0.22 | 0.11±0.18 | 0.085±0.14 | 0.074±0.12 | 0.063±0.097 | 0.054±0.095 | 0.057±0.10 | 0.057±0.092 | 0.069±0.12 | 0.041±0.083 |
| total age 0 recruit number | 8323.5±16732.8 | 7750.3±14896.6 | 5782.8±11599.4 | 3479.3±7535.8 | 6579.9±12886.2 | 5812.1±11200.5 | 8050.7±15081.8 | 3463.1±7271.0 | 4192.3±9077.9 | 3656.4±7152.8 |
| age 0 recruit sex ratio (proportion of female) | 0.57±0.22 | 0.56±0.19 | 0.48±0.13 | 0.52±0.22 | 0.47±0.20 | 0.44±0.18 | 0.38±0.19 | 0.48±0.23 | 0.55±0.17 | 0.52±0.25 |

**Table S2.** Summary of recovery period (30 years) model outputs (mean ± sd) from the shovelnose sturgeon spatially explicit individual-based model simulations for energetic (reduced energy allocation to female gonad development) and demographic (skewed sex ratio) perturbation scenarios. Vt_low, Vt_mid, and Vt_high indicate vitellogenin reduction in adult females exposed to low-, mid-, and high- level synthetic androgen. SS_low, SS_mid, and SS_high indicate skewed sex ratio in larvae exposed to low-, mid-, and high- level synthetic androgen.

| model output | baseline | Vt_low | Vt_mid | Vt_high | SS_low | SS_mid | SS_high | Vt + SS_low | Vt + SS_mid | Vt + SS_high |
| --- | --- | --- | --- | --- | --- | --- | --- | --- | --- | --- |
| **adult female** |  |  |  |  |  |  |  |  |  |  |
| sex ratio (proportion of female) | 0.31±0.11 | 0.31±0.09 | 0.31±0.09 | 0.31±0.08 | 0.31±0.11 | 0.29±0.09 | 0.28±0.13 | 0.30±0.09 | 0.28±0.09 | 0.29±0.11 |
| mean age of adult female | 6.69±0.17 | 6.71±0.23 | 6.67±0.21 | 6.51±0.64 | 6.70±0.31 | 6.67±0.24 | 6.80±0.47 | 6.64±0.28 | 6.67±0.20 | 6.55±0.13 |
| reproductive female abundance | 2282.5±1024.0 | 2958.4±1212.7 | 2044.6±831.4 | 837.8±426.5 | 2087.0±1034.9 | 822.3±404.0 | 1133.6±1052.6 | 1458.2±713.4 | 1867.9±1183.2 | 1022.3±516.7 |
| mean relative storage mass of reproductive female | 0.25±0.02 | 0.25±0.02 | 0.25±0.02 | 0.26±0.02 | 0.25±0.02 | 0.25±0.02 | 0.25±0.02 | 0.25±0.01 | 0.25±0.02 | 0.26±0.01 |
| mean relative storage mass of non-reproductive | 0.24±0.01 | 0.24±0.01 | 0.24±0.01 | 0.23±0.02 | 0.23±0.04 | 0.22±0.04 | 0.22±0.04 | 0.23±0.02 | 0.23±0.02 | 0.23±0.03 |
| mean relative gonad mass of reproductive female | 0.09±0.01 | 0.09±0.01 | 0.09±0.01 | 0.09±0.01 | 0.09±0.02 | 0.09±0.01 | 0.09±0.01 | 0.09±0.01 | 0.09±0.01 | 0.09±0.01 |
| number of spawner | 436.0±356.5 | 504.6±459.0 | 355.9±335.3 | 145.9±122.4 | 357.5±258.6 | 137.6±114.4 | 209.9±291.4 | 256.8±172.1 | 346.8±341.3 | 178.4±133.2 |
| mean spawning location (rkm) | 102.1±6.3 | 100.6±6.2 | 102.6±7.9 | 102.7±6.8 | 103.4±11.3 | 104.1±7.8 | 102.3±7.0 | 104.0±6.8 | 101.8±7.8 | 102.6±7.3 |
| mean spawning time (day of year) | 119.0±5.6 | 121.5±6.4 | 120.0±7.6 | 120.7±6.0 | 118.8±9.6 | 119.7±6.6 | 118.3±8.2 | 114.8±6.4 | 119.4±7.0 | 119.0±8.4 |
| mean egg number per female | 33886.9±1414.8 | 33499.4±1177.1 | 33925.7±1221.6 | 33833.5±1285.7 | 34562.2±1880.3 | 34199.4±1487.3 | 34551.2±1463.8 | 34768.0±1365.8 | 34104.0±1659.9 | 34905.2±1858.5 |
| mean spawning interval (years) | 3.18±0.36 | 3.08±0.43 | 3.24±0.42 | 3.21±0.47 | 3.27±0.38 | 3.24±0.36 | 3.09±0.37 | 3.31±0.31 | 3.18±0.36 | 3.22±0.40 |
| **early life stages** |  |  |  |  |  |  |  |  |  |  |
| total egg number (millions) | 14.7±12.1 | 17.2±11.9 | 12.1±9.03 | 4.87±3.99 | 12.1±8.48 | 4.71±3.94 | 8.39±12.4 | 8.85±5.88 | 11.8±11.6 | 5.90±5.35 |
| mean larval settlement location (rkm) | 96.8±8.9 | 95.9±10.0 | 97.6±10.9 | 94.6±9.5 | 97.7±10.8 | 99.0±11.2 | 98.2±13.7 | 99.1±10.3 | 97.6±9.0 | 95.9±10.8 |
| mean first-year survival rate (%) | 0.11±0.19 | 0.13±0.17 | 0.11±0.15 | 0.13±0.17 | 0.076±0.011 | 0.13±0.22 | 0.076±0.15 | 0.11±0.18 | 0.28±0.99 | 0.20±0.24 |
| total age 0 recruit number | 9340.3± 15274.4 | 11045.1±16848.3 | 6584.9±10952.2 | 3191.5±4420.9 | 5497.5±10621.3 | 2761.4±4475.5 | 2030.62±4863.7 | 3920.8±5213.8 | 7666.8±14721.8 | 4188.1±5734.9 |
| age 0 recruit sex ratio (proportion of female) | 0.56±0.16 | 0.52±0.10 | 0.51±0.11 | 0.51±0.16 | 0.51±0.20 | 0.53±0.18 | 0.52±0.22 | 0.42±0.17 | 0.47±0.17 | 0.53±0.23 |

## REFERENCES

Ancona, S., Dénes, F.V., Krüger, O., Székely, T. & Beissinger, S.R. (2017) Estimating adult sex ratios in nature. Philosophical Transactions of the Royal Society B: Biological Sciences, 372, 20160313.

Ankley, G.T., Jensen, K.M., Makynen, E.A., Kahl, M.D., Korte, J.J., Hornung, M.W., Henry, T.R., Denny, J.S., Leino, R.L. & Wilson, V.S. (2003) Effects of the androgenic growth promoter 17-β-trenbolone on fecundity and reproductive endocrinology of the fathead minnow. Environmental Toxicology and Chemistry, 22, 1350–1360.

Arcand Hoy, L.D. & Benson, W.H. (1998) Fish reproduction: an ecologically relevant indicator of endocrine disruption. Environmental Toxicology and Chemistry, 17, 49–57.

Arukwe, A. & Goksøyr, A. (1998) Xenobiotics, xenoestrogens and reproduction disturbances in fish. Sarsia, 83, 225–241.

Beddington, J. (1975) Mutual interference between parasites or predators and its effect on searching efficiency. Journal of Animal Ecology, 331–340.

Bernhardt, E.S., Rosi, E.J. & Gessner, M.O. (2017) Synthetic chemicals as agents of global change. Frontiers in Ecology and the Environment, 15, 84–90.

Braaten, P.J., Fuller, D.B., Holte, L.D., Lott, R.D., Viste, W., Brandt, T.F. & Legare, R.G. (2008) Drift dynamics of larval pallid sturgeon and shovelnose sturgeon in a natural side channel of the upper Missouri River, Montana. North American Journal of Fisheries Management, 28, 808–826.

Bramblett, R.G. & White, R.G. (2001) Habitat use and movements of pallid and shovelnose sturgeon in the Yellowstone and Missouri rivers in Montana and North Dakota. Transactions of the American Fisheries Society, 130, 1006–1025.

Coltman, D.W., O’Donoghue, P., Jorgenson, J.T., Hogg, J.T., Strobeck, C. & Festa-Bianchet, M. (2003) Undesirable evolutionary consequences of trophy hunting. Nature, 426, 655–658.

Crain, C.M., Kroeker, K. & Halpern, B.S. (2008) Interactive and cumulative effects of multiple human stressors in marine systems. Ecology Letters, 11, 1304–1315.

Dai, L., Vorselen, D., Korolev, K.S. & Gore, J. (2012) Generic indicators for loss of resilience before a tipping point leading to population collapse. science, 336, 1175–1177.

DeAngelis, D., Goldstein, R. & O’Neill, R. (1975) A model for tropic interaction. Ecology, 881–892.

del Carmen Alvarez, M. & Fuiman, L.A. (2005) Environmental levels of atrazine and its degradation products impair survival skills and growth of red drum larvae. Aquatic Toxicology, 74, 229–241.

Delonay, A.J., Chojnacki, K.A., Jacobson, R.B., Albers, J.L., Braaten, P.J., Bulliner, E.A., Elliott, C.M., Erwin, S.O., Fuller, D.B., Haas, J.D., Ladd, H.L.A., Mestl, G.E., Papoulias, D.M. & Wildhaber, M.L. (2016) Ecological requirements for pallid sturgeon reproduction and recruitment in the Missouri River—A synthesis of science, 2005 to 2012. Scientific Investigations Report, pp. 242. Reston, VA.

DeLonay, A.J., Jacobson, R.B., Papoulias, D.M., Simpkins, D.G., Wildhaber, M.L., Reuter, J.M., Bonnot, T.W., Chojnacki, K.A., Korschgen, C.E. & Mestl, G.E. (2009a) Ecological requirements for pallid sturgeon reproduction and recruitment in the Lower Missouri River: a research synthesis 2005-08.

DeLonay, A.J., Jacobson, R.B., Papoulias, D.M., Simpkins, D.G., Wildhaber, M.L., Reuter, J.M., Bonnot, T.W., Chojnacki, K.A., Korschgen, C.E., Mestl, G.E. & Mac, M.J. (2009b) Ecological requirements for pallid sturgeon reproduction and recruitment in the Lower Missouri River: A research synthesis 2005–08. U.S. Geological Survey Scientific Investigations Report 2009–5201.

Donald, P.F. (2007) Adult sex ratios in wild bird populations. Ibis, 149, 671–692.

Gascoigne, J., Berec, L., Gregory, S. & Courchamp, F. (2009) Dangerously few liaisons: a review of mate-finding Allee effects. Population Ecology, 51, 355–372.

Glebe, B. & Leggett, W. (1981) Temporal, intra-population differences in energy allocation and use by American shad (*Alosa sapidissima*) during the spawning migration. Canadian Journal of Fisheries and Aquatic Sciences, 38, 795–805.

Goto, D. (2009) Impacts of habitat degradation on Fundulus heteroclitus (Linnaeus) in urban tidal salt marshes in New York. Ph.D. Dissertation, City University of New York, New York NY.

Goto, D., Hamel, M.J., Hammen, J.J., Rugg, M.L., Pegg, M.A. & Forbes, V.E. (2015) Spatiotemporal variation in flow-dependent recruitment of long-lived riverine fish: Model development and evaluation. Ecological Modelling, 296, 79–92.

Goto, D., Hamel, M.J., Pegg, M.A., Hammen, J.J., Rugg, M.L. & Forbes, V.E. (2018) Spatially dynamic maternal control of migratory fish recruitment pulses triggered by shifting seasonal cues. Marine and Freshwater Research, 69, 942–961.

Goto, D. & Wallace, W.G. (2010) Bioenergetic responses of a benthic forage fish (Fundulus heteroclitus) to habitat degradation and altered prey community in polluted salt marshes. Canadian Journal of Fisheries and Aquatic Sciences, 67, 1566–1584.

Gregory, S.D., Bradshaw, C.J., Brook, B.W. & Courchamp, F. (2010) Limited evidence for the demographic Allee effect from numerous species across taxa. Ecology, 91, 2151–2161.

Grimm, V., Berger, U., DeAngelis, D.L., Polhill, J.G., Giske, J. & Railsback, S.F. (2010) The ODD protocol: a review and first update. Ecological Modelling, 221, 2760–2768.

Grüebler, M.U., Schuler, H., Müller, M., Spaar, R., Horch, P. & Naef-Daenzer, B. (2008) Female biased mortality caused by anthropogenic nest loss contributes to population decline and adult sex ratio of a meadow bird. Biological Conservation, 141, 3040–3049.

Gurney, W.S.C., Jones, W., Veitch, A.R. & Nisbet, R.M. (2003) Resource allocation, hyperphagia, and compensatory growth in juveniles. Ecology, 84, 2777–2787.

Hanson, P.C., Johnson, T.B., Schindler, D.E. & Kitchell, J.F. (1997) Fish bioenergetics 3.0. University of Wisconsin Sea Grant Institute, Madison, WI.

Hendry, A.P., Farrugia, T.J. & Kinnison, M.T. (2008) Human influences on rates of phenotypic change in wild animal populations. Molecular Ecology, 17, 20–29.

Holmes, E. & York, A. (2003) Using age structure to detect impacts on threatened populations: a case study with Steller sea lions. Conservation Biology, 17, 1794–1806.

Hope, B.K. (2005) Performing spatially and temporally explicit ecological exposure assessments involving multiple stressors. Human and Ecological Risk Assessment, 11, 539–565.

Houde, E.D. (1989) Comparative growth, mortality, and energetics of marine fish larvae: temperature and implied latitudinal effects. Fishery Bulletin, 87, 471–495.

Hrabik, R., Herzog, D., Ostendorf, D. & Petersen, M. (2007) Larvae provide first evidence of successful reproduction by pallid sturgeon, Scaphirhynchus albus, in the Mississippi River. Journal of Applied Ichthyology, 23, 436–443.

Huntzinger, T.L. & Ellis, M.J. (1993) Central Nebraska river basins, Nebraska. JAWRA Journal of the American Water Resources Association, 29, 533–574.

Huse, G. (1998) Sex-specific life history strategies in capelin (*Mallotus villosus*)? Canadian Journal of Fisheries and Aquatic Sciences, 55, 631–638.

Hutchings, J.A. (2015) Thresholds for impaired species recovery. Proceedings of the Royal Society B: Biological Sciences, 282, 20150654.

Jönsson, K.I. (1997) Capital and income breeding as alternative tactics of resource use in reproduction. Oikos, 57–66.

Kappenman, K.M., Webb, M.A.H. & Greenwood, M. (2013) The effect of temperature on embryo survival and development in pallid sturgeon *Scaphirhynchus albus* (Forbes & Richardson 1905) and shovelnose sturgeon *S. platorynchus* (Rafinesque, 1820). Journal of Applied Ichthyology, 29, 1193–1203.

Kjesbu, O.S., Solemdal, P., Bratland, P. & Fonn, M. (1996) Variation in annual egg production in individual captive Atlantic cod (*Gadus morhua*). Canadian Journal of Fisheries and Aquatic Sciences, 53, 610–620.

Kynard, B., Henyey, E. & Horgan, M. (2002) Ontogenetic behavior, migration, and social behavior of pallid sturgeon, Scaphirhynchus albus, and shovelnose sturgeon, S. platorynchus, with notes on the adaptive significance of body color. Environmental Biology of Fishes, 63, 389–403.

Lawson, D.M., Regan, H.M., Zedler, P.H. & Franklin, J. (2010) Cumulative effects of land use, altered fire regime and climate change on persistence of Ceanothus verrucosus, a rare, fire-dependent plant species. Global Change Biology, 16, 2518–2529.

Marr, A., Arcese, P., Hochachka, W., Reid, J.M. & Keller, L. (2006) Interactive effects of environmental stress and inbreeding on reproductive traits in a wild bird population. Journal of Animal Ecology, 75, 1406–1415.

Maxwell, S.M., Hazen, E.L., Bograd, S.J., Halpern, B.S., Breed, G.A., Nickel, B., Teutschel, N.M., Crowder, L.B., Benson, S. & Dutton, P.H. (2013) Cumulative human impacts on marine predators. Nature Communications, 4, 2688.

McBride, R.S., Somarakis, S., Fitzhugh, G.R., Albert, A., Yaragina, N.A., Wuenschel, M.J., Alonso-Fernández, A. & Basilone, G. (2015) Energy acquisition and allocation to egg production in relation to fish reproductive strategies. Fish and Fisheries, 16, 23–57.

Mills, L.J. & Chichester, C. (2005) Review of evidence: Are endocrine-disrupting chemicals in the aquatic environment impacting fish populations? Science of the Total Environment, 343, 1–34.

Molnar, P.K., Derocher, A.E., Lewis, M.A. & Taylor, M.K. (2007) Modelling the mating system of polar bears: a mechanistic approach to the Allee effect. Proceedings of the Royal Society B: Biological Sciences, 275, 217–226.

Morthorst, J.E., Holbech, H. & Bjerregaard, P. (2010) Trenbolone causes irreversible masculinization of zebrafish at environmentally relevant concentrations. Aquatic Toxicology, 98, 336–343.

Nash, J.P., Kime, D.E., Van der Ven, L.T.M., Wester, P.W., Brion, F., Maack, G., Stahlschmidt-Allner, P. & Tyler, C.R. (2004) Long-term exposure to environmental concentrations of the pharmaceutical ethynylestradiol causes reproductive failure in fish. Environmental Health Perspectives, 112, 1725.

NDEQ (2004) 2004 surface water quality integrated report. Water Quality Division. pp. 118 pp.

O’Brien, D., Lewis, S., Davis, A., Gallen, C., Smith, R., Turner, R., Warne, M., Turner, S., Caswell, S. & Mueller, J.F. (2016) Spatial and temporal variability in pesticide exposure downstream of a heavily irrigated cropping area: application of different monitoring techniques. Journal of Agricultural and Food Chemistry, 64, 3975–3989.

Pen, I., Uller, T., Feldmeyer, B., Harts, A., While, G.M. & Wapstra, E. (2010) Climate-driven population divergence in sex-determining systems. Nature, 468, 436.

Peters, E.J. & Parham, J.E. (2008) Ecology and management of sturgeon in the lower Platte River, Nebraska. Nebraska Technical Series.

Peterson, C.H., Rice, S.D., Short, J.W., Esler, D., Bodkin, J.L., Ballachey, B.E. & Irons, D.B. (2003) Long-term ecosystem response to the Exxon Valdez oil spill. Science, 302, 2082.

Rashleigh, B. & Grossman, G.D. (2005) An individual-based simulation model for mottled sculpin (Cottus bairdi) in a southern Appalachian stream. Ecological Modelling, 187, 247–258.

Rennie, M.D., Purchase, C.F., Lester, N., Collins, N.C., Shuter, B.J. & Abrams, P.A. (2008) Lazy males? Bioenergetic differences in energy acquisition and metabolism help to explain sexual size dimorphism in percids. Journal of Animal Ecology, 77, 916–926.

Ricker, W.E. (1975) Computation and interpretation of biological statistics for fish populations. Fisheries Research Board of Canada, Ottawa, Ontario.

Rideout, R.M., Rose, G.A. & Burton, M.P. (2005) Skipped spawning in female iteroparous fishes. Fish and Fisheries, 6, 50–72.

Scheffer, M., Baveco, J., DeAngelis, D., Rose, K. & Van Nes, E. (1995) Super-individuals a simple solution for modelling large populations on an individual basis. Ecological Modelling, 80, 161–170.

Stephens, P.A., Sutherland, W.J. & Freckleton, R.P. (1999) What is the Allee effect? Oikos, 185–190.

Takimoto, G. (2009) Early warning signals of demographic regime shifts in invading populations. Population Ecology, 51, 419–426.

Tilman, D. (1999) Global environmental impacts of agricultural expansion: the need for sustainable and efficient practices. Proceedings of the National Academy of Sciences, 96, 5995–6000.

Tilman, D., Fargione, J., Wolff, B., D’antonio, C., Dobson, A., Howarth, R., Schindler, D., Schlesinger, W.H., Simberloff, D. & Swackhamer, D. (2001) Forecasting agriculturally driven global environmental change. Science, 292, 281–284.

USDA (2001) USDA (U.S. Department of Agriculture, Agricultural Research) Nutrient Database for Standard Reference. http://www.nal.usda.gov/fnic/foodcomp.

Van Nes, E.H. & Scheffer, M. (2007) Slow recovery from perturbations as a generic indicator of a nearby catastrophic shift. The American Naturalist, 169, 738–747.

Wang, Y.L., Binkowski, F.P. & Doroshov, S.I. (1985) Effect of temperature on early development of white and lake sturgeon, *Acipenser transmontanus* and *A. fulvescens*. Environmental Biology of Fishes, 14, 43–50.

Wedekind, C. (2017) Demographic and genetic consequences of disturbed sex determination. Philosophical Transactions of the Royal Society B: Biological Sciences, 372, 20160326.

Weimerskirch, H., Lallemand, J. & Martin, J. (2005) Population sex ratio variation in a monogamous long-lived bird, the wandering albatross. Journal of Animal Ecology, 74, 285–291.

Welcomme, R.L. (1995) Relationships between fisheries and the integrity of river systems. Regulated Rivers: Research & Management, 11, 121–136.

Wildhaber, M., Papoulias, D., DeLonay, A., Tillitt, D., Bryan, J. & Annis, M. (2007a) Physical and hormonal examination of Missouri River shovelnose sturgeon reproductive stage: a reference guide. Journal of Applied Ichthyology, 23, 382–401.

Wildhaber, M.L., Papoulias, D.M., DeLonay, A.J., Tillitt, D.E., Bryan, J.L. & Annis, M.L. (2007b) Physical and hormonal examination of Missouri River shovelnose sturgeon reproductive stage: a reference guide. Journal of Applied Ichthyology, 23, 382–401.

Windsor, F.M., Ormerod, S.J. & Tyler, C.R. (2018) Endocrine disruption in aquatic systems: up-scaling research to address ecological consequences. Biological Reviews, 93, 626–641.

## REFERENCES FOR APPENDIX 1

Goto, D., Hamel, M.J., Hammen, J.J., Rugg, M.L., Pegg, M.A. & Forbes, V.E. (2015) Spatiotemporal variation in flow-dependent recruitment of long-lived riverine fish: Model development and evaluation. Ecological modelling, 296, 79–92.

Wildhaber, M.L., Papoulias, D.M., DeLonay, A.J., Tillitt, D.E., Bryan, J.L. & Annis, M.L. (2007) Physical and hormonal examination of Missouri River shovelnose sturgeon reproductive stage: a reference guide. Journal of Applied Ichthyology, 23, 382–401.

